# CYCLOPS: an open end-to-end platform for cyclic multiplex imaging and single-cell phenotyping

**DOI:** 10.64898/2026.06.11.731276

**Authors:** Sarwah Al-Khalidi, Nikki R. Paul, Mohammed Al-Khalidi, Ian Powley, Leo M. Carlin, Lucas Farndale, Ed Roberts

## Abstract

Multiplex immunofluorescent imaging enables deep spatial profiling of protein expression in tissues but is often limited by reliance on proprietary reagents, dedicated hardware, and closed analysis ecosystems. Here we present CYCLOPS (<u>Cycl</u>ic <u>O</u>pen <u>P</u>latform for <u>S</u>patial Proteomics), an end-to-end, open-source workflow for cyclic multiplex imaging and single-cell phenotyping using standard microscopy infrastructure. CYCLOPS integrates an Arduino-based automated fluidics system, an open-chamber stage insert, and antibody–oligonucleotide conjugation based entirely on published chemistries and off-the-shelf components. We demonstrate robust and reproducible antibody conjugation, high-quality multiplexed staining, and stable imaging across >10 cycles with minimal drift (<1 µm) and consistent fluorescence retention with low signal carry-over. The system supports efficient buffer exchange and consistent performance across multiple markers and imaging rounds. Using confocal microscopy, the workflow is compatible with three-dimensional imaging, enabling multiplexed analysis of volumetric tissue structures. To enable quantitative analysis, we establish an open-source image processing and analysis pipeline for single-cell feature extraction and phenotypic classification, avoiding reliance on proprietary software or black-box workflows. This framework integrates image registration, segmentation, and supervised classification to generate biologically interpretable single-cell data. Together, CYCLOPS provides a flexible and accessible platform for cyclic multiplex imaging, lowering barriers to adoption and enabling broader use of spatial proteomics across diverse research settings. This accessible framework democratizes high-plex imaging by enabling any laboratory with a standard confocal microscope to perform iterative multiplexing without reliance on proprietary reagents or hardware.

## INTRODUCTION

Spatially resolved, multiplexed immunofluorescent imaging (MIF) of proteins in tissues has transformed our ability to study cellular organization and interactions in complex biological systems^1^, with broad applications in immunology^2^, oncology^3^, and developmental biology^4^. Cyclic imaging approaches, which achieve high levels of multiplexing through iterative rounds of staining, imaging, and signal removal, have emerged as a powerful strategy for interrogating tissue architecture at single-cell resolution^5–7^. Several platforms for cyclic multiplex imaging have been developed, including co-detection by indexing (CODEX)^5^, iterative bleaching extends multiplexity (IBEX)^7^, and cyclic immunofluorescence (CycIF)^6^. These methods enable detection of tens to >50 protein markers within a single tissue section and have been widely applied to map complex tissue microenvironments. CODEX uses DNA-barcoded antibodies with iterative hybridization and imaging steps to achieve high multiplexing capacity^5^, whereas IBEX and CycIF rely on repeated cycles of staining and fluorophore bleaching or removal^6,7^.

Despite their power, these approaches present practical limitations that restrict broader adoption. Commercial implementations, particularly CODEX-like systems, although offering a ‘turn-key like’ experience, often rely on proprietary reagents, dedicated hardware, and closed software ecosystems, increasing cost and limiting experimental flexibility^8^. This also means images are obtained using a widefield microscope which can limit resolution in densely packed tissues and restricts to 2D imaging^9^.

Although IBEX and CycIF provide more open alternatives, they can require extensive manual handling, optimization of bleaching conditions, and careful panel design to maintain signal fidelity across cycles^6,7^. They are, however, compatible with confocal microscopes providing improved resolution, although this can introduce imaging artifacts when stitching large images^7^. In addition, cyclic imaging workflows are inherently sensitive to drift, signal carry-over, and cumulative tissue damage, necessitating robust experimental and computational correction strategies^6,7^.

Beyond data acquisition, analysis of multiplex imaging data remains a major challenge. Existing pipelines provide modular, open-source tools for image processing^10–12^, segmentation^13–15^, and quantification^16^, while specialized frameworks such as MuSpAn enable multiscale spatial analysis of cellular organization^17^. However, these approaches are often fragmented across multiple software environments and require substantial computational expertise. In contrast, some commercial platforms provide integrated analysis solutions implemented as black-box workflows, limiting transparency, reproducibility, and adaptability. As datasets increase in size and complexity, there is growing interest in integrating machine learning and artificial intelligence approaches for segmentation^13,14^, feature extraction^18^, and phenotypic classification^19^, further emphasizing the need for open and interoperable analysis pipelines.

To address these limitations, we developed CYCLOPS (<u>Cycl</u>ic <u>O</u>pen <u>P</u>latform for <u>S</u>patial Proteomics), an open and accessible cyclic imaging workflow that integrates hardware, chemistry, and analysis into a unified framework. CYCLOPS combines an Arduino-based automated fluidics controller^20^, adapted software^21^ to facilitate coordination of the fluidics with commercial confocal microscopy software, a universal open-chamber stage insert, and an antibody–oligonucleotide conjugation strategy derived entirely from published chemistries^22^ and off-the-shelf components. This approach eliminates dependence on proprietary reagents and hardware while maintaining compatibility with standard fluorescence microscopy. Importantly, the use of confocal microscopy enables extension of the workflow to three-dimensional imaging, allowing multiplexed analysis of volumetric tissue structures. We demonstrate that CYCLOPS enables stable imaging across repeated cycles, efficient and reproducible buffer exchange, and high-quality multiplexed staining using non-proprietary reagents. We also developed an approach to correct imaging artifacts introduced from stitching of large volume confocal images. In parallel, we establish an end-to-end image analysis pipeline built on open-source software, enabling transparent, reproducible, and accessible analysis of multiplex imaging data without reliance on proprietary or closed computational environments.

Here, we validate CYCLOPS through biochemical characterization of antibody–oligonucleotide conjugation, assessment of staining specificity and reproducibility, and quantitative evaluation of cyclic imaging performance. Together, this work provides an accessible and flexible framework for cyclic multiplex imaging and analysis, lowering barriers to adoption and enabling broader application of spatial proteomics across diverse research settings.

## RESULTS

### System Overview and Workflow

To establish an accessible cyclic multiplex imaging workflow, we integrated antibody–oligonucleotide conjugation, automated fluidics-driven staining, and quantitative image analysis into a unified platform (Fig. 1). Antibodies were conjugated to thiol-modified oligonucleotides using published chemistries^22^ and validated prior to use (Fig. 1A). Biological samples were embedded, sectioned, and stained with conjugated antibodies as described in Supplementary Protocol A (Fig. 1B).

**Figure 1:**
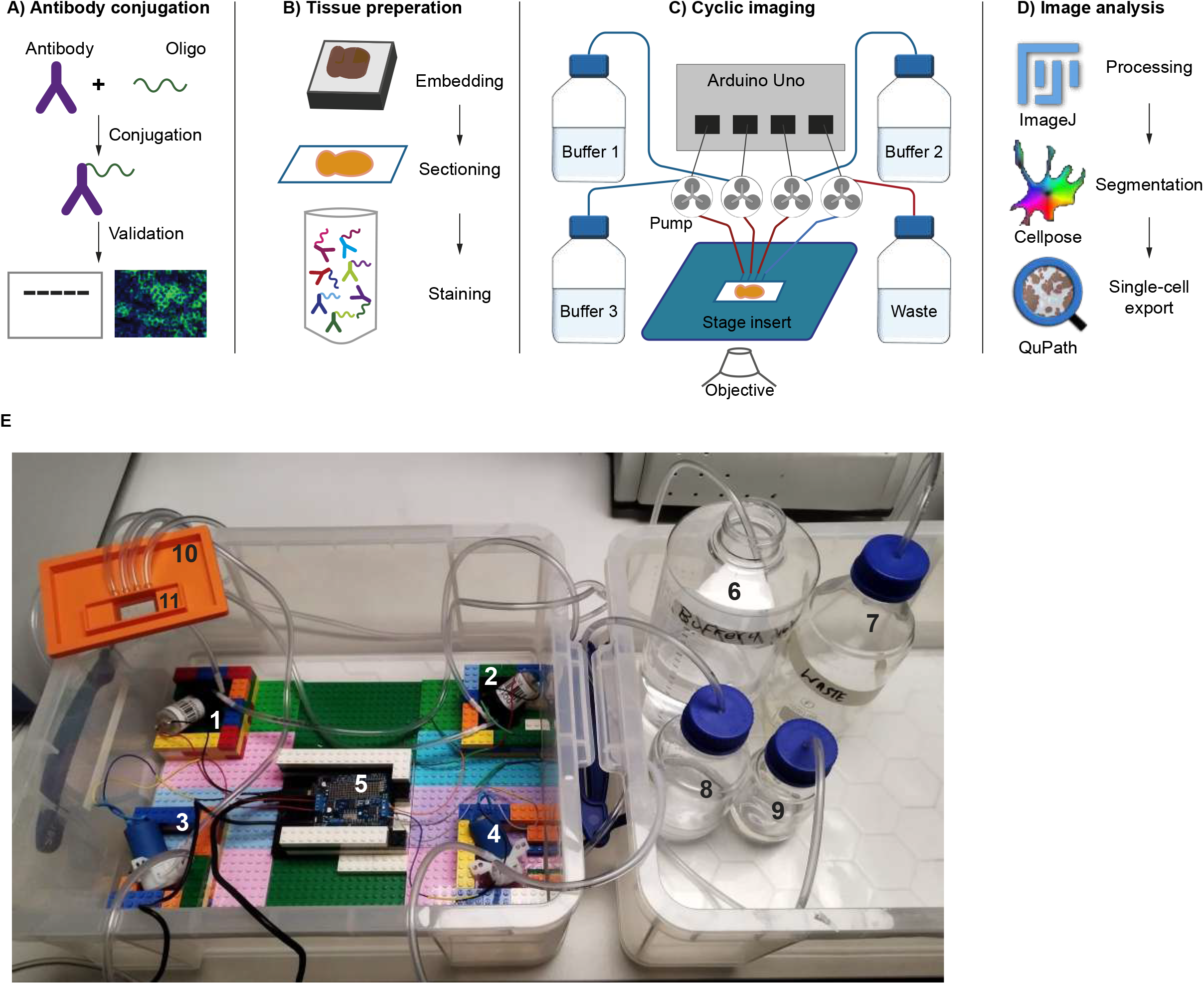
Overview of the open cyclic multiplex imaging workflow. **(A)** Antibody–oligonucleotide conjugation: Antibodies are coupled to thiol-modified DNA oligonucleotides using established crosslinking chemistries and validated prior to staining. **(B)** Tissue preparation: Fresh biological samples are embedded in OCT, sectioned, and stained with a master mix of conjugated antibodies using standard histological methods. **(C)** Automated cyclic imaging: An Arduino-controlled pump array connected to the imaging software delivers buffers from independent reservoirs to an open-chamber stage insert, enabling programmable hybridization, washing, imaging, and stripping cycles directly on a standard microscope stage. Waste is collected via an outlet reservoir. Blue lines, pump inlet. Red lines, pump outlet. **(D)** Image analysis: Acquired images are processed using open-source tools, including ImageJ for preprocessing, Cellpose for cell segmentation, and QuPath for single-cell feature extraction and export. **(E)** CYCLOPS cycler set up: Four pumps (1-4) are operated using an Arduino UNO chip and shield (5) programmed to pump consistent volume of buffers (6-9) into a microscope stage insert (10) fitted with a sample-mounted coverslip (11).

Automated cyclic imaging was enabled by a programmable microcontroller (Arduino Uno) controlling multiple independent pumps connected to dedicated buffer reservoirs (Fig. 1C, E). Buffers were delivered directly to an open-chamber stage insert 3D printed using a custom designed stage ([CYCLOPS/Hardware](https://github.com/alkhalidis/CYCLOPS/tree/main/Hardware)). The periphery of this stage can be adapted to fit different microscope stages, and the inner diameters can be altered to fit a variety of coverslip-mounted sample sizes, allowing sequential hybridization, washing, imaging, and stripping steps without removing the sample from the microscope. This configuration supports iterative staining cycles using non-proprietary reagents while maintaining compatibility with standard microscope stages.

Acquired images were processed using widely available open-source software tools, including ImageJ^11^ for preprocessing, Cellpose^13^ for segmentation, and QuPath^16^ for single-cell feature extraction (Fig. 1D). Together, this modular pipeline establishes a fully open framework for cyclic multiplexed imaging.

### Validation of Antibody–Oligonucleotide Conjugation

To generate DNA-tagged antibodies compatible with cyclic imaging using non-proprietary reagents, we evaluated multiple conjugation strategies based on published protocols. Initial testing of alternative conjugation approaches resulted in variable performance, with inconsistent recovery of functional antibody conjugates under our experimental conditions. We therefore adopted a maleimide–thiol conjugation strategy from a published method^22^, which yielded reproducible and efficient labeling across multiple antibodies. The protocol is well documented in the original publication and uses standard, commonly available reagents^22^. This protocol was also substantially cheaper than alternatives^5^ rendering this more accessible.

Using this approach, increasing the oligonucleotide-to-antibody ratio resulted in a progressive increase in A260/A280 values and higher molecular weight bands, consistent with higher oligonucleotide incorporation (Fig. 2A). This increase in labeling was accompanied by a corresponding enhancement in fluorescence signal following hybridization with complementary readout oligonucleotides (Fig. 2B). Based on these results, the higher conjugation condition (++) was selected for subsequent experiments. Under these conditions, conjugation produced the expected spectral shift, with increased absorbance in the 260 nm range compared to unconjugated antibodies (Fig. 2C), and a clear mobility shift on gel electrophoresis (Fig. 2D). These effects were consistent across multiple antibodies, with the degree of labelling showing low variability (coefficient of variation 10.7%) demonstrating reproducible conjugation efficiency (Fig. 2E).

**Figure 2:**
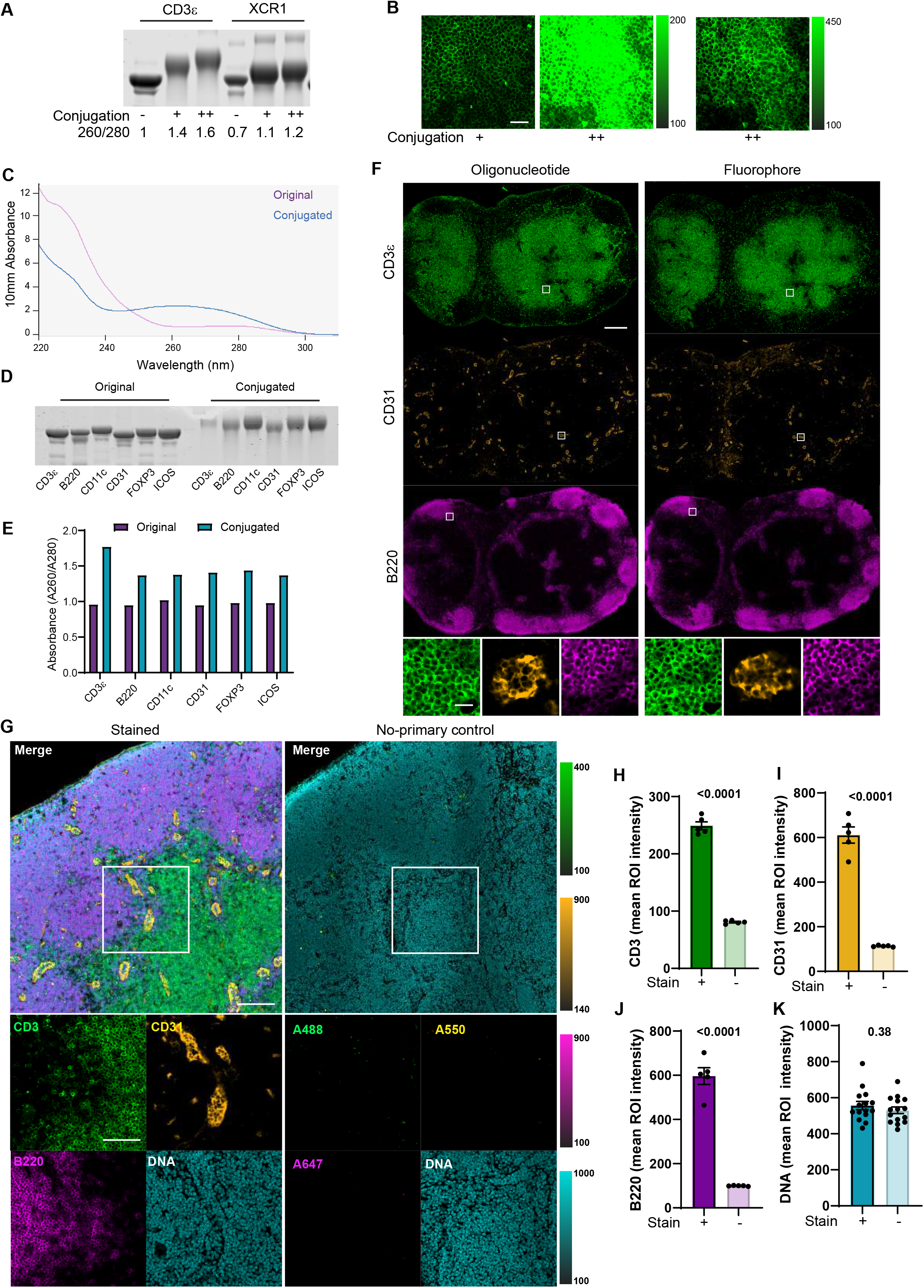
Validation of antibody–oligonucleotide conjugation and staining performance. **(A)** Antibody conjugation assessed by gel electrophoresis and A260/A280 ratios for selected antibodies before (−) and after low (+) and high (++) conjugation ratio, indicating increased nucleic acid content. **(B)** Representative fluorescence signal following hybridization with complementary oligonucleotides of low (+) and high (++) antibody labelling ratios. Far-right panel: high antibody labelling ratio visually adjusted for better assessment of staining quality. Scale bar = 50µm. **(C)** UV–visible absorbance spectra of unconjugated (purple) and conjugated (blue) antibodies. **(D)** Gel electrophoresis analysis demonstrating mobility shifts of conjugated antibodies relative to unconjugated controls (original). **(E)** Quantification of A260/A280 ratios across multiple antibodies before and after conjugation. **(F)** Comparison of staining using oligonucleotide-based detection versus directly fluorophore-conjugated antibodies for three markers. Scale bar = 250µm. Bottom panel: zoomed in images of boxed regions to show quality of staining. Scale bar = 20µm. **(G)** Representative tissue staining using conjugated antibodies (left) and no-primary antibody control (right). Scale bar = 100µm. Insets show higher magnification views of boxed regions. Scale bar = 50µm. **(H–K)** Quantification of mean fluorescence intensity (MFI) in regions of interest for the indicated markers, demonstrating significantly higher signal in stained samples compared to controls (-) (H-J) with no change in the nuclear signal (K). *n* = 5 ROIs. Statistical differences were determined by a t-test. Bars are means ± SEM.

To confirm that conjugation did not impair antibody functionality, we compared staining of mouse lymph node tissue obtained using oligonucleotide-based detection with that obtained using directly fluorophore-conjugated antibodies. Mouse lymph nodes were used as they consist of compartmentalized, thus easily distinguishable, cell subsets making comparisons more reliable. Both approaches yielded comparable staining patterns across three markers, indicating preserved antigen-binding specificity (Fig. 2F). We next assessed multiplex staining capability and specificity. Simultaneous staining with three oligonucleotide-conjugated antibodies followed by hybridization with complementary readout oligonucleotides produced distinct, non-overlapping signals corresponding to each target, demonstrating accurate marker co-registration and clear separation of cellular features (Fig. 2G). Expected anatomical staining patterns were preserved, and no detectable cross-reactivity or signal bleed-through was observed between oligos (Fig. 2G, SFig. 1), demonstrating the specificity and orthogonality of the oligo–antibody system in a multiplexed context.

To quantify staining quality, we measured signal-to-background ratios within the same images by comparing mean fluorescence intensity in tissue regions of positive staining to adjacent negative regions. All markers exhibited high signal-to-background ratios, indicating strong contrast and minimal non-specific background (Fig. 2H–K). Together, these results confirm that the conjugation and staining workflow yields high-quality images suitable for cyclic imaging.

### Cyclic Imaging Performance

We next evaluated the performance of the automated fluidics system during cyclic imaging. To assess spatial stability, images from each cycle were registered to Cycle 1 using translation transformations. Mean positional drift was 0.8 ± 0.3 µm across 10 cycles (Fig. 3A) and was effectively corrected using image registration, enabling accurate alignment of cells across cycles. To assess tissue and signal stability, we performed sequential rounds of staining and imaging using oligonucleotide-conjugated antibodies. Representative images acquired at early and late cycles showed consistent spatial patterns and signal intensity, with no detectable loss of staining fidelity (Fig. 3B). Negative control cycles confirmed efficient signal removal between rounds, with minimal residual fluorescence observed following stripping (Fig. 3B, lower panels).

**Figure 3:**
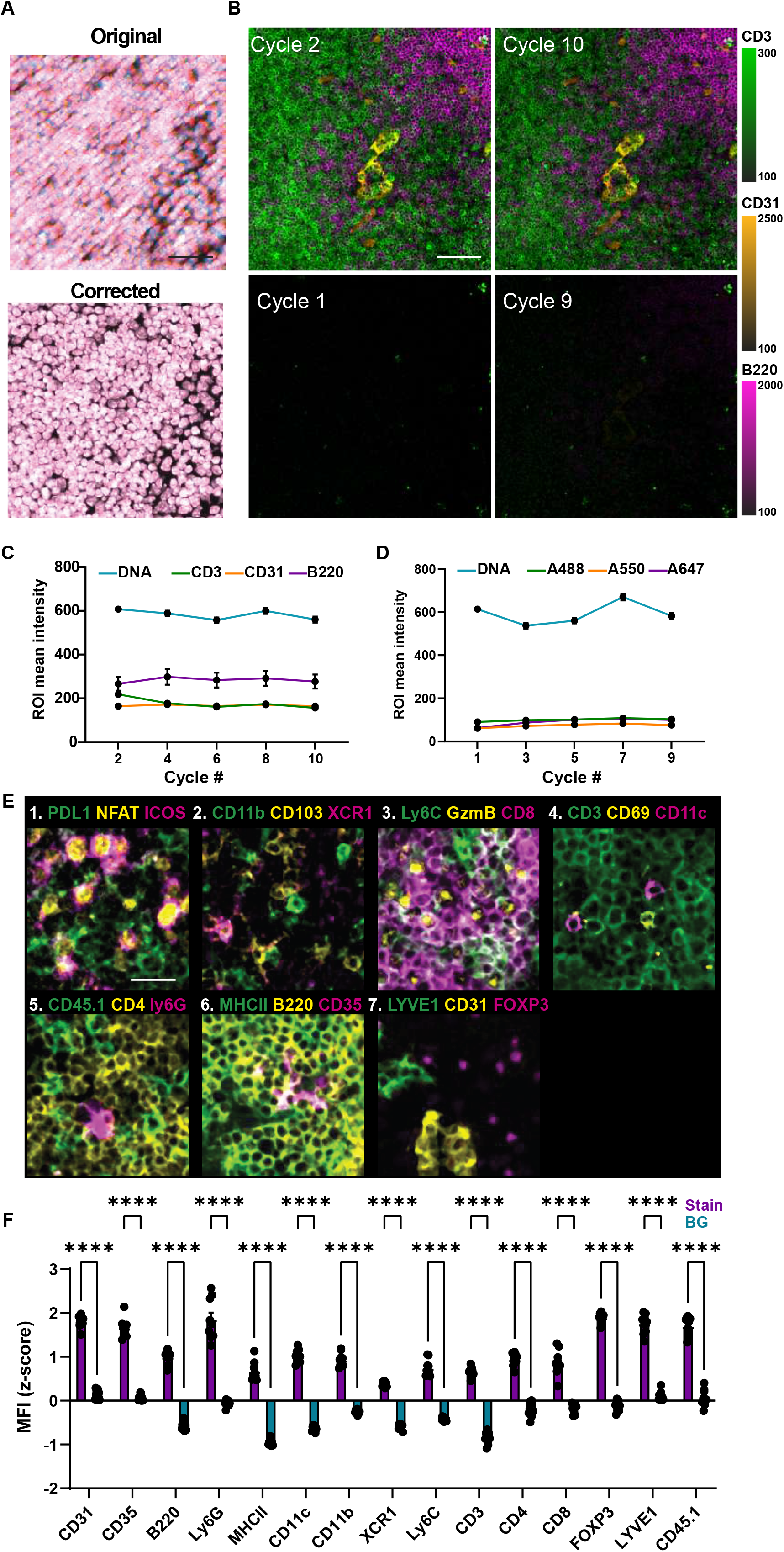
Performance of cyclic imaging workflow. **(A)** Spatial stability across imaging cycles. Maximum intensity projection of the nuclear channel across 10 cycles before (original, top) and after (corrected, bottom) cycle co-registration. Scale bar = 50µm. **(B)** Signal stability and stripping efficiency across cycles. Representative images from early and late cycles show consistent staining patterns and signal intensity. Lower panels show negative control cycles following stripping, demonstrating efficient removal of signal with minimal residual fluorescence. Scale bar = 50µm. **(C)** Quantification of fluorescence intensity retention across 10 imaging cycles. *n* = 9 ROIs. Bars are means ± SEM. **(D)** Background signal (cycle 1) and residual fluorescence retention following stripping. *n* = 9 ROIs. Statistical differences were determined by a paired t-test. Bars are means ± SEM. **(E)** Representative images showing distinct and biologically consistent staining patterns across >20 markers acquired over 7 cycles, with clear spatial separation between cell populations. Scale bar = 20µm. **(F)** Signal-to-background quantification for indicated markers. Mean fluorescence intensity (MFI) was measured in regions of positive staining (Stain) and compared to adjacent marker-negative regions (BG) within the same field of view, demonstrating strong signal-to-background separation.

Quantitative analysis of fluorescence intensity demonstrated stable signal levels for all markers, with only minor variation across up to 10 imaging cycles (Fig. 3C). Fluorescence intensities remained within 15% of initial values, while background levels after each wash were <10% of initial fluorescence (Fig. 3D), indicating minimal carry-over. Comparable results were observed across multiple detection channels, indicating consistent performance independent of fluorophore or target.

To demonstrate high multiplexing capability, we applied the workflow to a panel of markers spanning multiple cell types using the automated fluidics system to perform cyclic imaging, successfully acquiring >20 markers over 7 cycles. Distinct and biologically consistent staining patterns were observed for all targets, including immune and stromal markers, with clear spatial separation between cell populations (Fig. 3E). To assess the quality of staining using this system, the normalized mean fluorescent intensity of positively stained cells for each marker was compared to unstained cells in several independent biological samples, which confirmed strong separation between signal and background across all markers (Fig. 3F). Together, these results demonstrate that the system enables robust, reproducible cyclic imaging with minimal drift, stable signal retention, efficient signal removal, and high multiplexing capacity.

### Image Preprocessing and Illumination Correction

Because images are captured in tiles, stitching artifacts manifest as a repeating grid of darker regions. To account for grid-like stitching artifacts and illumination inconsistencies generated during image acquisition, a custom flat-field correction pipeline was applied to all raw images containing the artifact (Fig. 4A). The process relied on estimating the systematic imaging artifact from the tissue itself to ensure robust normalisation. To prevent background regions from skewing the illumination correction, an initial global tissue mask was generated using the nuclear channel of each cycle. As the unstained areas of the tissue are the same intensity as the non-tissue areas of background, naïve application of thresholding to obtain a tissue mask results in a grainy, inconsistent tissue mask that treats cytoplasm as background. To ensure that the tissue mask was internally smooth while allowing for holes, the nuclear image was down-sampled by a factor of 10 and smoothed with a 10×10 median filter to reduce high-frequency noise (Fig. 4B), before a global Otsu threshold was applied to this filtered image to separate tissue from background. To eliminate stray debris and imaging artifacts, connected component analysis was performed, and only the largest contiguous object was retained as the primary tissue mask of interest (Fig. 4C).

**Figure 4:**
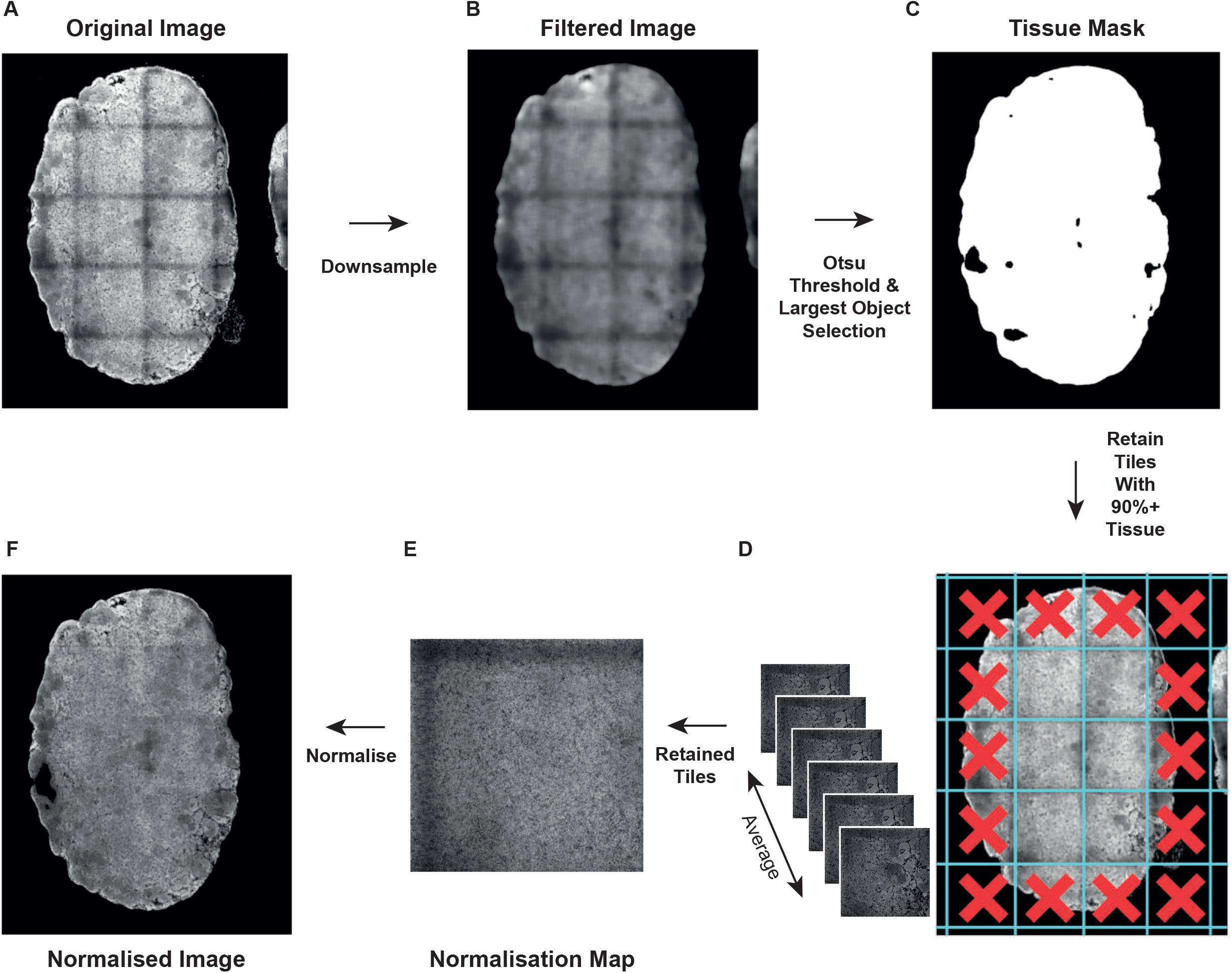
Image preprocessing and illumination correction pipeline. **(A)** Raw stitched image showing grid-like intensity artifacts arising from tile-based image acquisition. **(B)** Pre-processed nuclear image following down-sampling and median filtering to reduce high-frequency noise and improve robustness of tissue detection. **(C)** Tissue mask generated by global Otsu thresholding and connected component analysis, with retention of the largest contiguous object to define the primary tissue region. **(D)** Tile selection for illumination correction. The image was subdivided into patches corresponding to the microscope field of view, and only tiles containing ≥90% tissue (based on the mask in C) were retained for downstream processing. **(E)** Generation of a normalization map by averaging retained tiles to isolate systematic illumination artifacts while excluding biological structure. The resulting consensus patch is tiled to reconstruct a full-field correction map. **(F)** Illumination-corrected image obtained by dividing the raw image by the normalization map, resulting in removal of grid-like stitching artifacts and improved intensity uniformity across the tissue.

To isolate the microscope-specific stitching-related grid pattern of darker regions, the nuclear channel was divided into patches matching the microscope’s field of view (FOV) dimensions (e.g., 924×924 pixels). To ensure the normalization map was driven by biological signal rather than empty space, patches were filtered against the initial tissue mask; only patches containing at least 90% tissue were retained (Fig. 4D). The average pixel values across these patches were then taken to generate a single “consensus” patch that isolated the systematic dark-edge artifact without capturing specific tissue structures, yielding a normalisation map which could be applied across each FOV (Fig. 4E).

The single consensus patch was tiled sequentially to reconstruct a complete normalisation map matching the exact dimensions of the original image. As each channel is affected by this systematic artefact equally, all channels in the raw multiplexed image stack were then divided by this full-size normalisation map, successfully neutralising the grid-like stitching artifacts across the tissue (Fig. 4F).

### End-to-end image analysis pipeline for single-cell phenotypic classification

To enable quantitative interpretation of cyclic imaging data, we developed an end-to-end image analysis pipeline for single-cell feature extraction and phenotypic classification (Fig. 5). Raw image stacks from sequential imaging cycles were first co-registered in ImageJ to correct for positional drift, ensuring spatial alignment across all rounds (Fig. 5A). The aligned time-series data were then restructured by converting imaging cycles into channels, generating a unified multiplex image containing all measured markers. Redundant nuclear channels were removed, retaining a single reference channel for downstream analysis. Cell segmentation was performed in CellPose using the nuclear channel to generate segmentation masks, enabling accurate delineation of individual cells (Fig. 5B). These masks were superimposed on multichannel images imported into QuPath and used to extract per-cell fluorescence intensities across all markers, producing a high-dimensional single-cell dataset (Fig. 5C).

**Figure 5:**
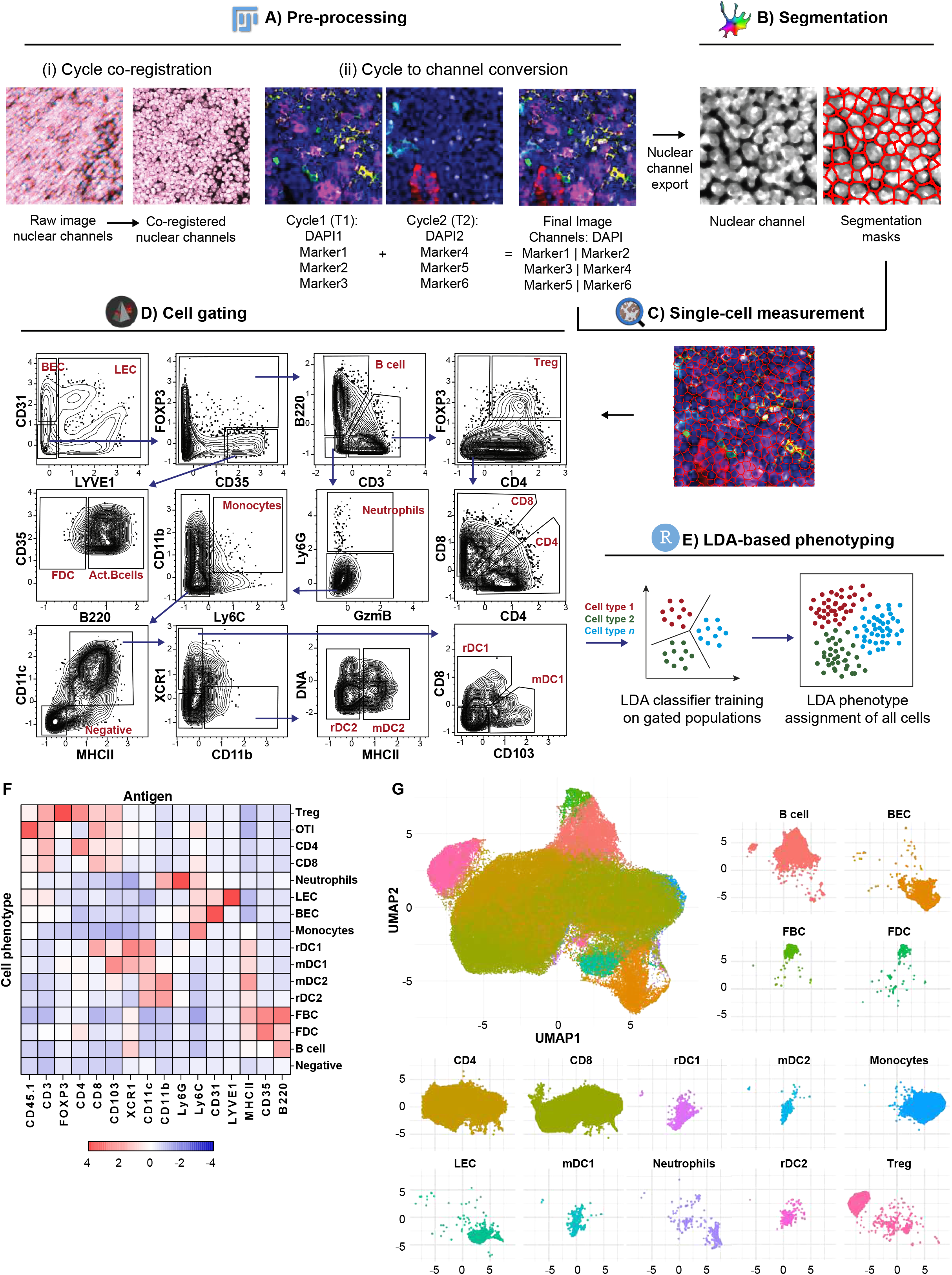
End-to-end image analysis pipeline for single-cell phenotypic classification. **(A)** Pre-processing of cyclic imaging data using ImageJ. Raw images from sequential imaging cycles were co-registered to correct for positional drift (I, reused form 3A). The aligned time-series data were then restructured by converting imaging cycles into channels, generating a unified multiplex image containing all markers (ii). Redundant nuclear channels were removed, retaining a single reference channel for downstream analysis. **(B)** Cell segmentation using Cellpose. The nuclear channel was used to generate segmentation masks, enabling delineation of individual cells. **(C)** Single-cell measurement acquisition using QuPath. Segmentation masks from Cellpose were applied to final images acquired from ImageJ to extract per-cell fluorescence intensities across all markers, producing a high-dimensional single-cell dataset. **(D)** Cell gating using FlowJo. Single-cell data were imported into FlowJo, where sequential gating strategies were applied to define reference cell populations based on marker expression. **(E)** LDA-based phenotyping. Gated populations were exported and used to train a linear discriminant analysis (LDA) classifier in R, which was subsequently applied to assign phenotypic labels to all cells in the dataset. **(F)** Heatmap of marker expression across annotated cell populations, showing expected marker enrichment patterns for each phenotype. **(G)** UMAP visualization of single-cell data colored by assigned phenotype, demonstrating clear separation of distinct cell populations.

To define phenotypic populations, single-cell data were imported into FlowJo, where sequential gating strategies were applied to identify reference populations based on marker expression (Fig. 5D). While this is not open source software, alternatives exist that can carry out flow cytometric gating which offer open source alternatives^23,24^. These gated populations were exported and used to train a supervised classification model using linear discriminant analysis (LDA) in R. The trained model was then applied to assign phenotypic labels to all cells in the dataset (Fig. 5E). This approach enabled consistent classification of diverse immune and stromal cell populations across multiplexed datasets. The resulting phenotypic assignments recapitulated expected marker expression patterns, as visualized by clustered heatmaps of marker intensity across cell types (Fig. 5F). In addition, dimensionality reduction using UMAP revealed clear separation of phenotypically distinct cell populations, further supporting the accuracy and consistency of the classification framework (Fig. 5G, SFig. 2). Together, this pipeline provides a scalable and reproducible framework for linking cyclic imaging data to biologically interpretable single-cell phenotypes.

## DISCUSSION

In this study, we present CYCLOPS (<u>Cycl</u>ic <u>O</u>pen <u>P</u>latform for <u>S</u>patial Proteomics), an integrated and accessible framework for cyclic multiplex imaging that combines open hardware, non-proprietary chemistry, and a fully transparent analysis pipeline. By unifying antibody–oligonucleotide conjugation, automated fluidics-driven staining, and single-cell analysis, CYCLOPS provides a complete end-to-end workflow for spatial proteomics using standard microscopy infrastructure. Our results demonstrate that this approach achieves robust and reproducible performance across all stages of the workflow. Antibody–oligonucleotide conjugation yielded consistent labeling efficiency across multiple antibodies, with preserved staining specificity and strong signal-to-background ratios. The fluidics-driven cyclic imaging system enabled stable multi-round imaging, with minimal positional drift and low signal carry-over across up to 10 cycles. Together, these results indicate that non-proprietary reagents and open hardware can support high-quality multiplex imaging comparable to more complex and integrated systems.

A key advantage of CYCLOPS is its flexibility relative to existing cyclic imaging platforms. Commercial systems, particularly CODEX-like implementations^8^, provide highly optimized and automated workflows but rely on proprietary reagents, dedicated hardware, and closed software ecosystems. While IBEX and CycIF offer more open alternatives, they typically require extensive manual intervention and optimization of bleaching or stripping conditions^6,7^. In contrast, CYCLOPS decouples cyclic imaging from vendor-specific constraints, enabling users to adapt conjugation chemistries, modify staining panels, and integrate the workflow with existing microscopy platforms. This flexibility is particularly valuable for method development, custom panel design, and applications where access to commercial systems is limited.

In addition to experimental flexibility, CYCLOPS emphasizes transparency and reproducibility in data analysis. By integrating widely used open-source tools for image processing, segmentation, and feature extraction, and combining these with supervised phenotypic classification, the pipeline avoids reliance on proprietary or “black-box” analysis environments. This approach enables users to interrogate and adapt each stage of the analysis workflow and facilitates integration with downstream spatial analysis frameworks such as MuSpAn for multiscale characterization of tissue organization. As spatial proteomics datasets continue to grow in complexity, this openness will be increasingly important for incorporating emerging machine learning and artificial intelligence approaches for segmentation, feature extraction, and phenotypic classification.

The modular design of CYCLOPS further enhances its adaptability. Individual components of the workflow—including the fluidics system, stage insert, conjugation strategy, and analysis pipeline—can be independently modified or extended. This enables incorporation of alternative chemistries, expanded marker panels, or improved computational methods without requiring changes to the overall system architecture. Such modularity provides a foundation for community-driven development and iterative refinement of both experimental and computational components.

Despite these advantages, several limitations should be considered. The current implementation requires initial setup and optimization of hardware and conjugation protocols, which may introduce variability between laboratories. While we demonstrate stable imaging across multiple cycles and markers, further validation across a wider range of tissue types, imaging systems, and experimental conditions will be important to establish generalizability. Furthermore, while using confocal imaging improves resolution and provides 3D imaging option, the time required for imaging is longer than that required on widefield microscopes. In addition, although the workflow reduces dependence on proprietary systems, it does not yet match the level of automation and throughput offered by fully integrated commercial platforms. Future work could focus on further standardization, automation of sample handling, and benchmarking against established methods.

An important feature of this framework is its compatibility with confocal microscopy, which enables extension to three-dimensional imaging. While the present study focuses on two-dimensional tissue sections, the ability to acquire optical sections across depth provides a clear path toward multiplexed volumetric imaging of intact tissues. This capability could enable analysis of cellular organization and interactions in three dimensions, further enhancing the biological insight obtained from cyclic imaging datasets. Beyond technical performance, CYCLOPS addresses a critical barrier to adoption in spatial proteomics: accessibility. The use of off-the-shelf components for the fluidics system and non-proprietary reagents for antibody conjugation substantially reduces both upfront and per-experiment costs compared to commercial platforms. This lowers the barrier to entry for laboratories without access to dedicated instrumentation and enables broader dissemination of multiplex imaging approaches.

Together, this work establishes CYCLOPS as a flexible and accessible platform for cyclic multiplex imaging and analysis. By integrating open hardware, chemistry, and software into a unified workflow, this approach lowers barriers to adoption while maintaining robust performance. As spatially resolved single-cell analysis continues to expand, such open and modular frameworks will play an important role in enabling widespread application and continued methodological innovation.

## Supporting information

Protocols

## Acknowledgments

We thank P. Thomason and C. Mitchell, (CRUK Scotland Institute) as well as L. Lemgruber Soares and S. Baillie (University of Glasgow) for providing technical support and assistance in data acquisition. We would like to thank the core services and advanced technologies at the CRUK Scotland Institute (RRID:SCR_027384) with particular thanks to the Biological Services Unit and the Beatson Advanced Imaging Resource (RRID:SCR_023875). We also thank Catherine Winchester (CRUK Scotland Institute) for critically reviewing this manuscript.

## Funding

This work was supported by CRUK core funding to the Scotland Institute (A31287) and CRUK core funding to E.W.R. (A1920) and to L.M.C. (A23983). Work was also supported by a PCUK programme funding to E.W.R. (MA-TIA22-001).

## Author contributions

S.A.K.: Conceptualization, Methodology, Software, Validation, Formal analysis, Data curation, Investigation, Visualisation, Writing draft, Project administration. N.P.: Methodology, Software. L.F., I.P.: Software. L.M.C.: Methodology. E.R.: Conceptualization, Methodology, Investigation, Writing draft, Visualisation, Supervision, Project administration, Funding acquisition. All authors critically reviewed the manuscript.

## Declaration of Interests

L.M.C. has consulted for Ono Pharmaceuticals UK and is an academic mentor to Biomed X Heidelberg on unrelated work. All other authors declare that they have no competing interests.

## Data and code availability

All data needed to evaluate the conclusions in the paper are present in the paper or the Supplementary Materials. All code used for automated fluidics control, cyclic imaging, image preprocessing, and single-cell analysis is available at [CYCLOPS](https://github.com/alkhalidis/CYCLOPS). This includes Arduino scripts for pump control, image processing workflows implemented in ImageJ, and analysis pipelines for segmentation, feature extraction, and phenotypic classification in R. Example configuration files and a demonstration dataset are provided to facilitate reproducibility.

## Figure legends

**Supplementary Figure 1: Specificity of oligonucleotide–antibody detection and minimal cross-reactivity**

Tissue sections were stained with a single oligonucleotide-conjugated primary antibody per condition (CD3, CD31, or B220), followed by simultaneous incubation with three complementary fluorescent readout oligonucleotides (A488, A555, and A647). For each antibody, fluorescence signal was observed exclusively in the corresponding detection channel, with minimal to no signal detected in non-matching channels, demonstrating high specificity of oligonucleotide hybridization and minimal cross-reactivity between probes. Representative images show the nuclear (DNA) channel, individual fluorescence channels, and merged images for each condition. These results confirm the orthogonality of the oligonucleotide–antibody system and support its suitability for multiplexed cyclic imaging. Intensity scales are shown for each channel. Scale bar = 50µm.

**Supplementary Figure 2: Marker expression across UMAP embedding of single-cell data**

UMAP representation of single-cell data generated from cyclic imaging, showing expression levels of individual markers projected onto the same embedding as shown in Fig. 5. Each panel displays the normalized expression of a single marker, with color intensity indicating relative expression levels across cells. The distribution of marker expression supports the phenotypic classification and cell population assignments described in Fig. 5.

**Supplementary Figure 3: Arduino-based fluidics control system**

**(A)** Arduino Uno microcontroller with motor driver shield used for automated control of peristaltic pumps. Key components are indicated, including solder points for wiring, power jumper, external power input, USB interface, and pump connection terminals (P1-P4).

**(B)** Representative peristaltic pump used for buffer delivery, with tubing connections and polarity of electrical wiring indicated. Pumps are connected to the Arduino-controlled driver board to enable programmable delivery of reagents during cyclic imaging, where the other end of the wire soldered to the positive terminal of the pump (+) is attached to the positive terminal on the Arduino shown in A, and the wire soldered to the negative terminal of the pump (-) is attached to the negative terminal of the Arduino.

**(C)** Final set up of the stage insert. The sample-mounted coverslip is adhered to the insert using a gasket, and the tubes are fitted into the stage inlets, with the waste outlet sitting at a lower level compared to buffer inlets. The insert is then mounted on the microscope stage and secured in place using BlueTack (not shown, indicated by blue lines) to ensure minimal image drift.

## METHODS

### Animal tissues

All mice were maintained under specific pathogen-free conditions and treated in accordance with the Animal Welfare and Ethical Review Body Committee at the University of Glasow in compliance with the UK Home Office regulations (ASPA, 1986, PPL P72BA642F). All mice were bred and housed at the CRUK Scotland Institute.

### Antibody–Oligonucleotide conjugation

Antibodies were conjugated to thiol-modified DNA oligonucleotides using a maleimide–thiol coupling strategy adapted from published protocols. Oligonucleotide sequences were designed based on previously published CODEX barcoding strategies^5^. Antibody–oligonucleotide conjugation was performed using a maleimide–thiol coupling protocol adapted from Immuno-SABER staining methods^22^. Briefly, 5′-thiol-modified oligonucleotides (Integrated DNA Technologies) were reduced with 100mM dithiothreitol (DTT) for 2h at room temperature in the dark and purified using NAP-5 desalting columns (GE Healthcare) to remove excess reducing agent. Antibodies formulated in PBS were concentrated to ∼2mgml^−1^ using 50kDa molecular weight cut-off filters (Amicon Ultra, Millipore) and reacted with maleimide–PEG2–succinimidyl ester crosslinker (Thermo Fisher Scientific) for 1.5h at 4°C. Excess crosslinker was removed using Zeba desalting columns (7kDa cutoff). Activated oligonucleotides were incubated with antibodies at defined molar ratios (low (+) and high (++) conjugation conditions) overnight at 4°C. Conjugated antibodies were washed six times using 50kDa centrifugal filters to remove unbound oligonucleotides and stored at 4°C until use.

Successful conjugation was assessed by measuring absorbance at 260 and 280 nm using a Nanodrop spectrophotometer. A260/A280 ratios were compared between unconjugated and conjugated antibodies. UV–Vis absorbance spectra (200–350 nm) were recorded to confirm increased nucleic acid absorbance. Conjugates were further validated by gel electrophoresis, where mobility shifts relative to unconjugated antibodies were used as an indicator of successful modification. Full conjugation and validation protocol used was published by Saka et.al^22^. Functional validation was performed by staining tissue sections using oligonucleotide-conjugated antibodies followed by hybridization with complementary fluorescent oligonucleotides, and comparing staining patterns to those obtained using directly fluorophore-conjugated antibodies, as described below.

### Tissue preparation and staining

Mouse lymph nodes were harvested in PBS, embedded and frozen in optimal cutting temperature (OCT) freezing media, sectioned, and mounted onto coverslips using standard histological procedures. Prior to staining, sections were briefly defrosted, fixed in acetone for 10min, air-dried, rehydrated, then fixed in 1.6% paraformaldehyde (PFA). After blocking at room temperature for 30min, samples were incubated overnight at 4°C with oligonucleotide-conjugated antibodies diluted in block buffer using optimised concentrations, then washed thoroughly to remove unbound antibodies. Multiplex staining was performed using panels of conjugated antibodies, followed by hybridization with complementary fluorescent readout oligonucleotides. Full tissue preparation and staining protocols were adapted from published protocols^5^ and can be found in Supplementary Protocol A. Antibody and oligo details can be found in Supplementary Table 1.

### Fluidics device construction and stage insert fabrication

An automated fluidics system was constructed using an Arduino Uno microcontroller connected to four independent peristaltic pumps. Each pump line connects a buffer reservoir to a dedicated inlet leading to a custom open-chamber stage insert. Stage inserts were 3D printed from acrylic to fit standard microscope stages. The stage insert consists of a rectangular open cavity that accommodates a glass coverslip to allow imaging via an inverted microscope. Four angled inlet ports deliver buffer directly to the chamber. Full device construction details can be found in Supplementary Protocol B, and stage 3D printing files are provided in [CYCLOPS/Hardware](https://github.com/alkhalidis/CYCLOPS/tree/main/Hardware).

Fluidics operation was controlled via custom scripts adapted from prior fluidic controller scripts^21^ enabling automated execution of cyclic staining, washing, and stripping steps. Buffer exchange was achieved by sequential activation of pumps, delivering defined volumes of reagents directly into the chamber while waste was removed via an outlet channel. Details of the buffer exchange steps can be found in Protocol B, and the device control script can be found in [CYCLOPS/Cyclic_Imaging](https://github.com/alkhalidis/CYCLOPS/tree/main/Cyclic%20Imaging).

### Cyclic imaging workflow

Coverslips mounted with stained tissue sections were sealed within the stage insert chamber using an elastomeric gasket and removable sealant. The stage insert was fixed on a Nikon AXR inverted microscope for imaging using the 20x objective. Buffer exchange cycles were performed automatically according to predefined pump scripts run through the NIS software (6.10.02). For each round, cyclic imaging was performed by iteratively applying fluorescent readout oligonucleotides, imaging, and removing signal between cycles. Each cycle consisted of hybridization of spectrally separate fluorescent oligonucleotides tagged with AF488, ATTO555 or AF647 as well as a nuclear marker, washing to remove unbound probes, confocal xyz image acquisition using 20x magnification, then signal removal using stripping buffer. Imaging was performed using a standard confocal fluorescence microscope (Nikon AXR) with fixed acquisition settings across cycles using the 405nm, 488nm, 561nm and 640nm laser lines, with an XY pixel size of 0.43µm, an optical resolution of 0.44µm and an averaging of 2 and a 0.075 frames per second imaging time. Full cyclic setup details can be found in Supplementary Protocol C, and cyclic imaging script are provided in [CYCLOPS/Cyclic_Imaging](https://github.com/alkhalidis/CYCLOPS/tree/main/Cyclic%20Imaging). Antibody and oligonucleotide details used can be found in Supplementary Table 1.

### Image pre-processing

Stitched image stacks (xyztc) were imported into Fiji (v1.45f). Maximum intensity projection was first applied to all channels in all cycles, generating xytc images. Cycles were then co-registered by means of the nuclear channel of a reference cycle using translation-based StackReg registration to correct for positional drift. Following registration, image stacks were restructured by converting timepoints (cycles) into channels, generating a unified xyc multiplex image. A single nuclear channel was generated by applying minimum intensity projection to a stack of nuclear channels from all cycles, retaining a single reference channel for downstream analysis. The Fiji preprocessing code is provided in [CYCLOPS/ImageProcessing](https://github.com/alkhalidis/CYCLOPS/tree/main/ImageProcessing), and full implementation details are found in Supplementary protocol D.

### Cell segmentation and feature extraction

Cell segmentation was performed using the nuclear channel. Segmentation masks were generated using Cellpose (v2.0). Nuclear masks were expanded to approximate whole-cell boundaries where appropriate. Extracted masks were then superimposed on pre-processed images imported to QuPath (v0.4), and fluorescence intensities for all markers were extracted on a per-cell basis, generating a high-dimensional single-cell dataset. QuPath code for mask importation and single-cell intensity measurements is provided in [CYCLOPS/ImageProcessing](https://github.com/alkhalidis/CYCLOPS/tree/main/ImageProcessing), and full implementation details are found in Supplementary protocol D.

### Flow cytometry–style gating

CSV files of single-cell data exported from QuPath were imported into RStudio (2023.12.1+402) for normalization and data cleaning, where unique cell IDs were assigned. CSV files of single-cell data for each sample were then exported and imported into FlowJo (Treestar) software (V10.10.0) for manual gating. Sequential gating strategies were applied to define reference cell populations based on marker expression profiles. Gated populations corresponding to distinct cell types were exported as CSV files for downstream analysis.

### Phenotypic classification using LDA

Gated cell populations were used as training data for supervised classification using linear discriminant analysis (LDA) implemented in R. The trained LDA model was applied to the full single-cell dataset to assign phenotypic labels to all cells. Classification results were used for downstream visualization and analysis. LDA classification was validated through dimensionality reduction performed using UMAP to visualize high-dimensional single-cell data. Cells were colored according to assigned phenotypes to assess separation between populations. Mean marker expression across phenotypically defined populations was visualized using heatmaps for further validation.

### Quantification of signal, background and drift

Signal-to-background ratios were calculated by measuring mean fluorescence intensity (MFI) in regions of positive staining and comparing to adjacent marker-negative regions within the same field of view. Signal stability across cycles was assessed by tracking fluorescence intensity of a region of interest over repeated imaging rounds. Background carry-over was quantified as residual signal following stripping relative to initial fluorescence. Positional drift was quantified by measuring displacement of nuclear features across cycles following registration to a reference cycle. Mean drift was calculated in the x and y directions across multiple fields of view.

