## Supplementary material for "CYCLOPS: an open end-to-end platform for cyclic multiplex imaging and single-cell phenotyping": Protocols

**Supplementary Protocol A**

This protocol details the optimised procedure for sample freezing, sectioning and staining for multiplexed cyclic imaging, adapted from published protocols[5].

**Tissue preparation**

**Materials:**

- 1x phosphate buffered saline (PBS)
- Tissue-Tek® O.C.T. Compound (Labtech, 4583)
- Casset
- Methanol (VWR, 67-56-1)
- Dry ice
- Glass coverslips (Merck, CLS2980243)
- 0.1% poly-L-Lysine (Merck, P8920)

**Procedure:**

1. Harvest samples in PBS.
2. Embed sample in OCT:
    a. Roll sample on a piece of tissue paper.
    b. Put a small amount of OCT at the base of a size-appropriate cassette.
    c. Place the sample in the cassette and cover in OCT.
    d. Place a flat-bottom reservoir containing a small volume of methanol on dry ice. Place pieces of dry ice in the methanol until they no longer dissipate.
    e. Slowly place the cassette containing OCT-covered tissue in the methanol bath, ensuring methanol does not seep into the cassette as that will prevent OCT from solidifying.
    f. Once the OCT turns completely white, wait for a further 5min then store tissue at -80°C until sectioning.
3. Incubate 1.5mm coverslips in ready-made solution of poly-L-Lysine overnight at room temperature.
4. Wash with dd$H_2O$ and air dry.
5. Cut 10μm sections and mount on coverslips. Store in -80°C until staining.

**Tissue staining**

**Materials:**

- 1x PBS
- BSA
- 500mM EDTA pH 8.0
- Tris pH 8.0
- 5M NaCl
- Mouse IgG (Sigma, I5381)
- Rat IgG (Sigma)
- Sheared salmon sperm DNA, 10 mg/ml in $H_2O$ (ThermoFisher, AM9680).
- Blocking oligonucleotides (Integrated DNA Technology)
- 16% Paraformaldehyde
- BS3 crosslinker (Thermo Fisher Scientific, 21580)
- Acetone
- Methanol
- DMSO ampoules (Sigma, 1034240005)

**Buffers:**

- **TE buffer:** 10 mM Tris pH 8.0 (Teknova), 1 mM EDTA in $ddH_2O$.
- **Buffer1:** 0.5% BSA, 5mM EDTA in PBS. (100ml of PBS, 0.5g BSA, 1ml of 500mM EDTA).
- **Buffer2:** 200mM phosphate buffer, 0.5M NaCl in $dH_2O$, then diluted 1:1 in Buffer1.
- **Blocking reagent 1**: 2mg/ml mouse IgG in **Buffer2**.
- **Blocking reagent 2**: 2mg/ml rat IgG in **Buffer2**.
- **Blocking oligonucleotide**: 0.5mM in TE buffer.
- **Block Buffer:** 50µg/ml mouse IgG, 50µg/ml rat IgG, 0.5mg/ml sheared salmon sperm DNA, 3.5µM of each blocking oligonucleotide.
- **Buffer3:** 1:9 5M NaCl: Buffer1.
- **BS3 stock fixative solution:** 200mg/ml BS3 in DMSO from a freshly opened ampoule; stored at -20°C in 15µl aliquots.
- **Buffer 4:** 10mM Tris pH7.5, 10mM MgCl2, 0.1% Triton, 150mM NaCl.
- **Rendering buffer:** 20% DMSO (v/v) in buffer 4.
- **Rendering mix:** Fluorescent oligo (1:1000), Hoechst (1:500), Salmon sperm DNA (1:20) in rendering buffer.
- **Stripping buffer:** 80% DMSO (v/v) in buffer 4.

**Procedure:**

1. Defrost the samples at room temperature for 2min and dry by wiping the back of the coverslip with tissue paper.
2. Dip dried coverslip-mounted sections for 10min into room temperature acetone, then fully dry for 2min at RT.
3. Draw around the edges of the coverslip with a pap pen.
4. Rehydrate sections by incubating them in a Buffer1 bath for 5min. Reuse the Buffer1 bath for subsequent washes.
5. Place the coverslip in a humidity chamber. Unless otherwise stated, all incubation steps are carried out at room temperature in the humidity chamber using 100µl of solution.
6. Fix sections by covering them with an appropriate volume of Buffer1 supplemented with 1.6% paraformaldehyde (PFA) for 10 min at room temperature.
7. Wash PFA off with two changes of Buffer1.
8. Incubate in Buffer2 for 10min. Decant and flip off excess buffer, do not wash.
9. Incubate sections in block buffer for 30min.
10. Prepare staining master mix by adding conjugated antibodies at the desired final concentration in block buffer.
11. Decant sections and flip off excess buffer, do not wash.
12. Add the staining master mix to the coverslip and incubate overnight at 4°C in the humidity chamber with slow shaking.
13. Wash sections with three changes of Buffer2, 2min each.
14. Fix in 1.6% PFA diluted in Buffer3 for 10min.
15. Wash twice in a PBS bath to remove BSA, which will precipitate in methanol.
16. Incubate sections for 5min in ice cold methanol (methanol stored in -20°C).
17. Wash twice in a PBS bath. Leave in the second PBS bath till the next step is ready.
18. Defrost a 15µl 200mg/ml BS3 vial, add 985µL of PBS and mix well.
19. Incubate the tissue with BS3 in PBS (3mg per mL) for 20min.
20. Wash sections in a PBS bath then in a bath containing Buffer3.

**Tip:** Fixed specimen can be stored in Buffer3 at 4°C until the rendering multicycle experiment. This is stable for up to two weeks.

For single cycle imaging:

1. Wash sections in Buffer4.
2. Wash sections twice with rendering buffer, 5min each.
3. Add rendering mix and incubate for 5min.
4. Wash 3x in rendering buffer.
5. Wash 1x with Buffer4.
6. Mount in Buffer4 and image.

**Supplementary Protocol B**

This protocol consists of steps to assemble the cycler, which consists of two parts: the chip and the pumps. The chip is prepared is prepared by soldering the shield on to an Arduino chip. The pumps are prepared by soldering them to jumper wires. The cycler is then assembled by attaching the pump wires to the terminal blocks on the shield.

**Martials:**

| Item | Source | Code |
|---|---|---|
| 1x Arduino Uno Rev3 | PiHut | A000066 |
| 1x Adafruit Shield for Arduino v3 Kit | PiHut | ADA1438 |
| 1x Electric Soldering Iron | ScrewFix | 40326 |
| 1x USB-A to USB-B Cable | PiHut | 105562 |
| 1x 12V 2A Power Supply, 2.1mm Barrel Jack | PiHut | 106673 |
| 4x Peristaltic Liquid Pump 12V DC Power | PiHut | ADA1150 |
| Premium Male/Male Jumper Wires | PiHut | 103192 |
| Tygon® E-3603 Laboratory Tubing | Cole-Parmer | |

**Shield-Arduino UNO assembly:**

1. Break apart a 0.1" header into 6, 8 and 10-pin long pieces.
2. Slip the long ends into the headers of the Arduino.
3. Place the assembled shield on top of the Arduino so that all the short parts of the header are sticking through the outer set of pads.
4. Solder each one of the pins into the shield to make a secure connection.

**Pump assembly:**

1. Strip one end of two jumper wires to expose the metal centre.
2. Solder the exposed end of each of the wires to one terminal of Pump1 (P1).
3. Insert the other ends of the wires in the M1 terminal of the shield, noting the polarity of the connections as shown in SFig. 3A-B. Using a screwdriver, secure the connection by screwing the wire in place.
4. Repeat the process for Pump2, Pump3 and Pump4, inserting the wires in the M2, M3 and M4 terminals of the shield, respectively.
5. Attach the power jumper in place, as shown in SFig. 3a.
6. Remove the tubes that come with the pump, and insert an equal length of tube in the white connector of the pump
7. Attach the power and USB-B cables.

**Software installation**

This section contains code adapted from NanoJ-Fluidics GUI and enables users to install and operate the pump control software from within the NIS software as a macro. For other options please refer to the NanoJ-Fluidics GUI page.

1. Install the Arduino IDE

2. Install the Adafruit Motorshield Library through the Arduino IDE user interface

3. To operate the pumps, download the “1_Pump_Control_Software.mac” from CYCLOPS/Cyclic Imaging and place it in the Macro folder of the NIS system. This is commonly located in “C:\Program Files\Nikon\Shared\Macros” but user specific system may be different.

Alternatively, open the “1_Pump_Control_Software.mac” file as a text document, select all and copy, then go to the “Macro” tab within the NIS software, select “New”, paste the code and save it as “1_Pump_Control_Software.mac”.

4. Repeat step 3 for the “2_PreImage.mac and “3_PostImage.mac” files found in the same GitHub directory.

**Supplementary Protocol C**

This protocol contains details to carry out cyclic imaging using the CYCLOPS cycler assembled in Supplementary Protocol B and the Nikon AXR confocal microscope equipped with the *NIS*-Elements AR 6.10.02 software.

**Cycler setup:**

1. Switch on the PC and the microscope.
2. Connect the cycler to the PC using the USB-A cable, then connect the power cable to a power source and switch it on.
3. To fill the tubes and test the pumps, place the inlet tube of each pump in the flask containing its corresponding buffer, and place the outlet tube in the waste container. For the waste pump, both tubes are placed in the waste container.
4. Open the NIS software and wait for it to fully load.
5. Go to “Macro” drop down menu, select “run macro from file”
6. Select “1_Pump_Control_Software.mac”. A window will open with Pump 1 automatically selected (shown as P1.1). Wait for “Connected!” to appear.
7. To fill the cycler tubes, click “start”. This will start the first pump. Allow the buffer to completely fill the tube.
8. Once the buffer starts dripping into the waste container, wait for 10s then click stop.
9. Repeat steps 7 and 8 for the other two buffer pumps by selecting them from the drop-down menu, one at a time.
   **Tip:** You can also check that the waste pump is working by selecting it form the drop-down menu. Ensure that the inlet tube is not submerged in buffer, click start, allow it to tun for a few seconds then click stop.

Congratulations! You have successfully set up the cycler.

**Stage setup:**

1. Thoroughly clean the coverslip bed. Any dust, water or grease will prevent proper adhesion of the adhesive gasket, allowing buffer to seep onto the microscope.
2. Remove the ‘open-window’ side of the adhesive gasket cover. This will allow holding the exposed gasket without touching the adhesive layer.
3. Apply the gasket to the coverslip bed. Apply pressure to all sides of the gasket repeatedly till it is completely sealed.
4. Remove the other side of the gasket cover by gently flicking out one of its corners.
   **Tip 1:** make sure not to damage the gasket corners when removing the cover
   **Tip 2:** if the gasket comes off when removing the cover, remove the gasket completely and apply a fresh one. This could be due to remnants of grease from the previous experiment.
5. Dry the back of the coverslip thoroughly. any liquid will prevent proper adhesion to the gasket.
6. Place the coverslip on the gasket mounted in the coverslip bed. Apply pressure to all sides of the coverslip repeatedly till it is completely sealed to the gasket.
7. Invert the stage insert and apply grease to all adhesion points to seal the coverslip-gasket-insert trio together.
   **Tip:** Be careful not to apply grease in the imaging area of the coverslip. This will prevent the sample form being imaged and may stick to the objective.
   **Tip 2:** Be quick so that the sample is not left dry for longer than needed, but do not rush as ensuring proper adhesion will save a lot of pain in the long run.
8. Invert the stage insert back and apply buffer 4 to the sample, filling the coverslip chamber (~2ml). Let the stage sit for 5min on a paper towel to ensure no leaking is present.
9. Remove the filled outlet tubes from the waste container and connect each outlet tube of the cycler to one of the inlets on the stage insert.

10. Set the microscope objective to its lowest limit
11. Place the insert on the microscope stage and secure it in place using Bostik Blu Tack.
    **Tip:** It is important to ensure that stability of the insert to minimise movement during cyclic imaging, reducing image registration correction downstream.

**Cyclic imaging:**

For our study, cyclic imaging was done using the AXR confocal microscope, with a 20x objective and sequential acquisition using the 405nm, 488nm, 561nm and 640nm excitation lasers, with a pinhole size of 1AU at the 640nm laser, a zoom Size = 2, and an optical resolution = 0.44µm. These settings can be modified according to the user's requirement. The region of interest to be acquired was manually drawn during acquisition as described below, after which the xy dimensions of each image was determined by the software. The stack size was determined for each run depending on the flatness of the stage/insert/coverslip by taking a test stack before running the JOB.

For best practice, as the laser settings cannot be changed between cycles, excitation and acquisition settings for each laser should be adjusted during panel optimisation such that the range allows for the dimmest markers to be clearly distinguishable form background without saturation of the brightest markers to be acquired. These settings should be saved and included in the JOB to be used for all cycles.

1. With the coverslip-mounted stage secure on the microscope, prepare 1ml of rendering buffer with nuclear marker only
2. Navigate to "Run Macro From File" in the Macro drop down menu and run "2_PreImage". Add the 1ml of nuclear stain during the 5min incubation period. This is important to set the objective focus correctly, and the fluid level should be the same as the remining imaging cycles.
3. Ensure the objective is in the centre of the coverslip, then slowly move the objective towards the coverslip till it focusses on the tissue, being careful not to hit the coverslip or the grease surrounding it.
4. With the Z level at the centre of the tissue, click 'find perfect focus'. This will allow the microscope to calculate the distance between the objective and the coverslip, switching the PFS function and maintaining focus throughout imaging.
5. In the NIS software, navigate to "Analysis Controls" from the "View" drop down menu and select "JOBS Explorer".
6. In the JOBS Explorer window, import the "CYCLOPS_JOB.bin" script from CYCLOPS/Cyclic Imaging.
7. Switch on the DAPI channel so you can see the tissue down the eyepiece
8. In the JOBS menu of the NIS system, double click on the "Cyclic Imaging" script. This will activate it and allow you to go through all the steps to set the following:
    a. Select the microscope lasers imaging parameters. These are the parameters optimised during panel optimisation.
    b. Specify the number of cycles required. This depends on the number of markers to be imaged.
    c. Specify the number of Z stacks required. This depends on the flatness of the stage/insert/coverslip mounted.
    d. Register the upper left and bottom right corners of the tissue for the overview image.
9. Click start. The microscope will first produce an overview image.
10. Select the polygon tool, then on the overview image produced draw regions of interest (ROIs).
    **Tip:** ROIs will be numbered sequentially and can be renamed by right-clicking on them. Alternatively, a screenshot of the ROIs can be taken and the ROIs renamed downstream after image acquisition.
11. Once all the ROIs are drawn, go back to the JOBs screen and click "Continue".
12. The "PreImage.mac" macro will start:
    a. Withdraw liquid

b. Add rendering buffer
c. Wait for 60s
d. Withdraw liquid
**e. Wait for 5min – this is the incubation step for the secondary oligo, where 1ml of the secondary oligo mix should be added using a pipette.**
f. Withdraw liquid
g. Wash twice with rendering buffer, 1min each
h. Add Buffer4
i. Image

13. Once all ROIs have been imaged, the "PostImage.mac" macro will automatically start:
    a. Withdraw liquid
    b. Add stripping buffer 3x, 3min each.
    c. Wash 2x with rendering mix
    d. Add buffer4
    e. Wait for user to start next cycle
14. Click "Continue". This will start the next cycle, repeating steps 9 and 10 till all cycles specified in the JOBS have completed
15. Once all cycles are completed, the images will be saved by ROIs
16. Open each ROI in the NIS system, apply maximum intensity projection on Z, then stitch using "blending" on the MIP image. Both these options can be found in the "Processing" drop down menu in the "Image" tab.
17. Save the MIP stitched images for preprocessing in ImageJ.

**Supplementary protocol D**

**Cellsegmentation and single-cell data generation**

Software:

1. ImageJ:
    a. Install the latest stable version of FIJI.
    b. Install the HyperStackReg plugin
2. Cellpose: install cellpose and its dependencies. In our system, we installed Cellpose with Conda and operated it within the GPU as described in the Cellpose github
3. Install QuPath

Process:

1. In a new folder, create the following subfolders
    a. **1_raw_images**: place MIP stitched images
    b. **2_final_images**: place ImageJ output images
    c. **3_nuclear**: place nuclear channels exported from the ImageJ script to use for Cellpose segmentation
    d. **4_Project**: empty folder for QuPath
    e. **5_analysis:** empty folder to save single-cell data
    f. **Macros**: place the “CYCLOPS_ImageJ.ijm” script from Github
    g. Create a text file and name it “**channel_names.txt**”. Open the file empty file, type “Label”, hit enter, then list all channel names in order of imaging, each in a new line. Save the file and close.
2. Open ImageJ and load the CYCLOPS_ImageJ.ijm script by dragging it into the software.
3. Edit the script to insert the directory of the **channel_names.txt** file
4. Change the values of “channels” as described in the script file
5. Run the script, this will prompt choosing the **1_raw_images, 2_final_images** and **3_nuckear** files successively.
6. Once the script run is complete, check you should have final images and nuclear images in their appropriate folders
7. Load Cellpose. Drag and drop one of the nuclear images and train a segmentation script as described on the Cellpose github.
8. Save the resulting masks as a .tiff file in the **3_nuckear folder**, where “name of the mask file” = “name of nuclear image” masks.tiff
9. In QuPath, create a new project directed to the **4_Project** folder.
10. Load all images in **2_final_images** to the project.
11. Download the “importCellposeMaskAndMeasure.groovy” file from github and place it in a new folder named “scripts” created within the **4_Project** folder.
12. In QuPath, go to  and run the script after changing the path to reflect the folder where the masks are saved.
13. Save single-cell data to **5_analysis** folder by exporting measurements for detections from the “measure” menu. This will generate a .csv file of cells x measurement, which can be used for all downstream analysis.

**Cell LDA phenotyping**

Software:

1. Install R and RStudio
2. Install FlowJo

Process:

1. In the **5_anaysis** folder, now containing the “measurements.csv” file:
    a. Download the “Data_cleaning” and “LDA” R script from github and save it to this folder
    b. Create a **ForFlowJo** folder
2. Run the Data_cleaning script, which should generate .csv files for each image in the **ForFlowJo** folder.
3. Load the generated csv files into flowJo, gate on the populations of interest, then export each population for all samples as a csv file such that 5 populations of interest should generate 5 files.
4. Run the LDA script after adjusting the populations of interest.
5. Save the phenotypes generated as a csv file for further data analysis.
