## Supplementary figures and images for "CYCLOPS: an open end-to-end platform for cyclic multiplex imaging and single-cell phenotyping"

### Supplemental Figures

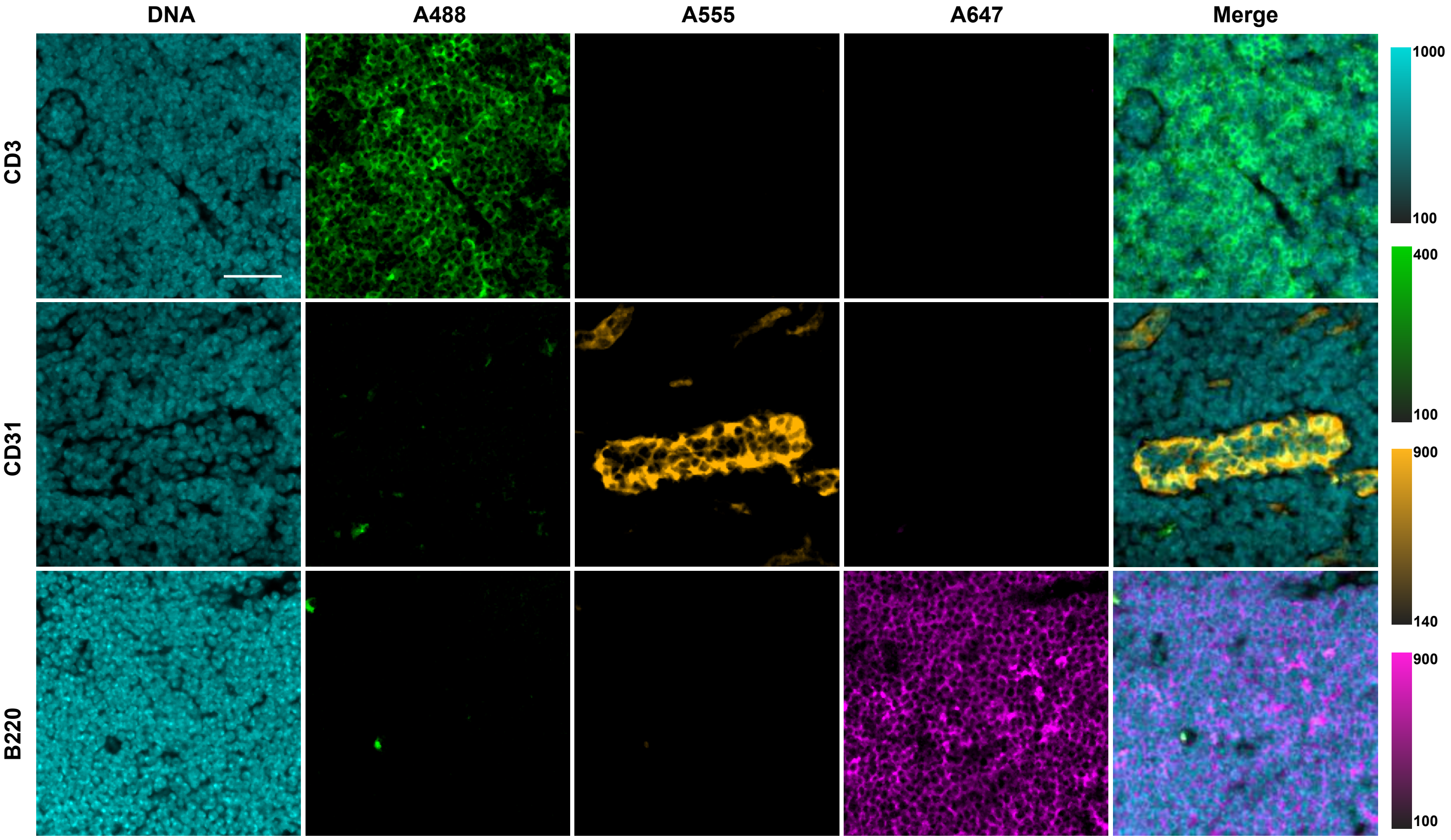

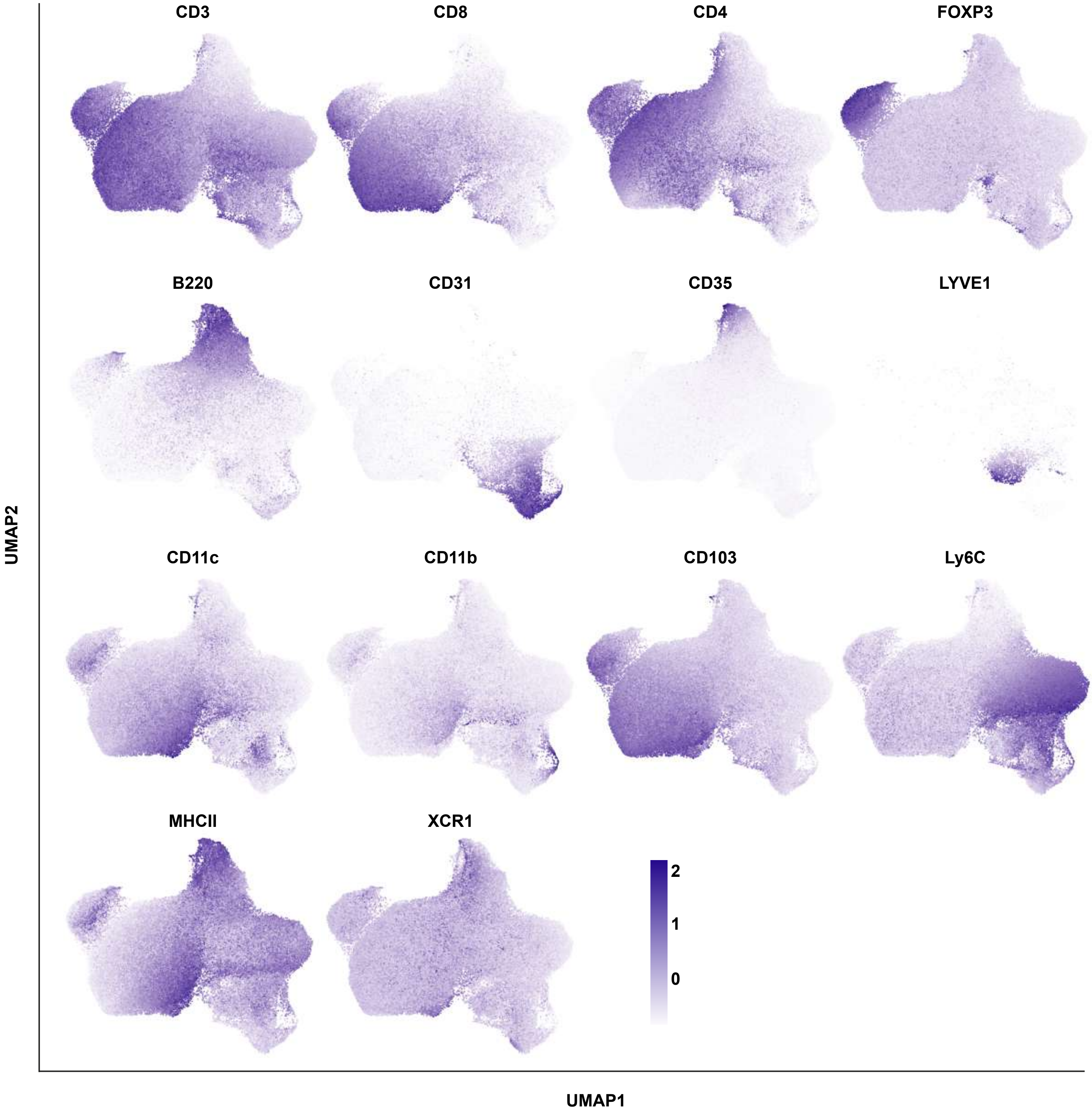

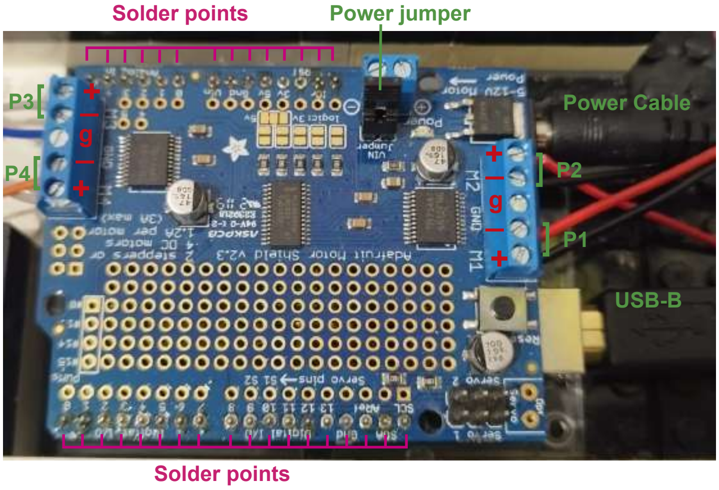

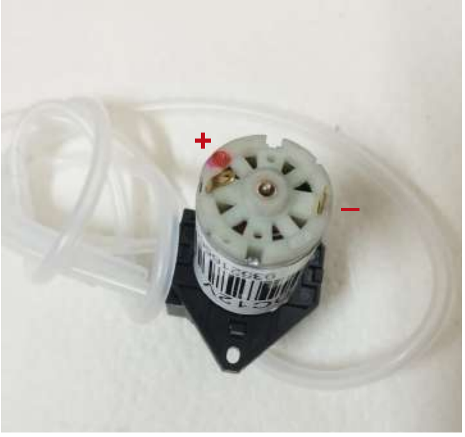

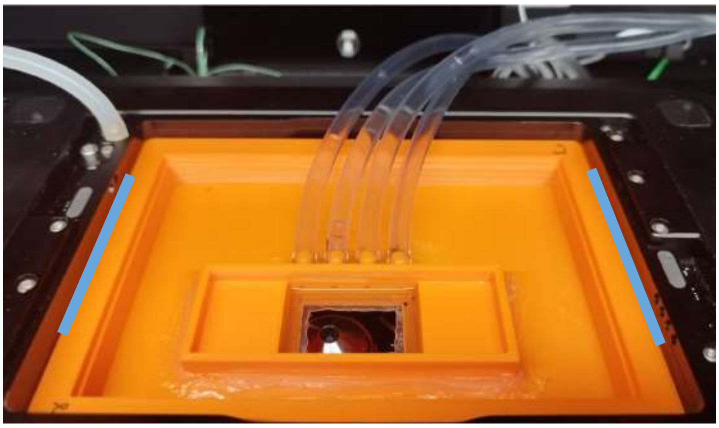
